# Complex versus simple models: ion-channel cardiac toxicity prediction

**DOI:** 10.1101/206946

**Authors:** Hitesh. B. Mistry

## Abstract

There is growing interest in applying detailed mathematical models of the heart for ion-channel related cardiac toxicity prediction. However, a debate as to whether such complex models are required exists. Here an assessment in the predictive performance between two established cardiac models, *gold-standard* and *cardiac safety simulator*, and a simple linear model *B_net_* was conducted. Three ion-channel data-sets were extracted from literature. Each compound was designated a cardiac risk category based on information within CredibleMeds. The predictive performance of each model within each data-set was assessed via a leave-one-out cross validation. In two of the data-sets *B_net_* performed equally as well as the leading cardiac model, *cardiac safety simulator*, both of these outperformed the *gold-standard* model. In the 3^rd^ data-set, which contained the most detailed ion-channel pharmacology, *B_net_* outperformed both cardiac models. These results highlight the importance of benchmarking models but also encourage the development of simple models.

## Introduction

There is a growing belief within the pharmaceutical industry that in order to improve predictions of future experiments more detailed mathematical models of biology are required.^1,2^ However by including more detail not only does the number of parameters that need to be estimated increase but so does the degree of structural uncertainty i.e. the degree of confidence in the actual structure of the equations.^3^ The objective of this study is to look at this issue within the field of drug induced ion-channel cardiac toxicity. This area has a well-defined question relating to prediction where a debate about the complexity of the model needed is ongoing.

The question of interest is: can high-throughput ion-channel screening data predict the propensity for a given type of arrhythmia, torsades de pointes (TdeP), in humans.^4^ In order to answer this question the literature is divided in terms of the complexity of the modelling approach required.^5^ The complex models used are based on biophysical models which describe the changes in ionic currents over time within a single cardiac cell.^6^ They contain 100s of parameters and 10s of differential equations. The drug input into these models involves scaling ion-channel conductance’s by the amount of block at a given drug concentration.^7^ Two biophysical models that have gained favour in the literature are the *gold-standard*, as described by Zhou *et al.*^8^, model by O’Hara and Rudy^9^ which is being put forward for use by regulatory agencies^10^ and another, by TenTusscher *et al.*^11^, forms a key part of the *cardiac safety simulator*.^12^ An alternative simpler model being put forward analyses the net difference, *via* a linear combination, in drug block of the ion-channels of interest, termed *B_net_*.^5^ Thus it is based on a higher level of abstraction than biophysical models and focusses on known biology/pharmacology.

Two previous studies have shown that the simple model is likely to give similar predictive performance to the more complex models.^5,13^ However in those studies the definition of torsadegenic risk lacked consistency as each data-set used different criterion. Furthermore those studies were based only on 3 ion-channels, hERG, Cav 1.2 and Nav 1.5 peak, and so the dimensionality of ion-channel space can be considered narrow.

In this study we analyse the predictive performance of the *gold standard*, *cardiac safety simulator* and *B_net_* models using a consistent and reliable definition of torsadegenic risk from CredibleMeds^14,15^ across three literature data-sets.^16–18^ Two of these data-sets, Mirams *et al.*^17^ and Kramer *et al.*^16^, measured drug effect against 3 ion-channels, hERG, Cav 1.2 and Nav 1.5 peak. The third and latest data-set, from Crumb *et al.* ^18^, considers drug effect on 7 ion-channels, hERG (IKr), KCNQ1 + KCNE1 (IKs), Kv4.3 (Ito), Kir2.1 (IK1), Cav 1.2 (ICaL), Nav1.5 peak (INa) and Nav1.5 late (INaL), the largest number studied so far.

By using a consistent definition of torsdagenic risk across different data-sets the analysis conducted will provide a detailed view on the performance of each model. Thus enabling scientists to make a more informed decision about which modelling approach is likely to be the most useful for the prediction problem considered.

## Methods

### Data

Ion-channel IC50 values, defined as concentration of drug the reduces the flow of current by 50%, were collected from three publications.^16–18^ Compounds within those data-sets were classified as being TdeP positive or TdeP negative based on their classification by Credible Meds.^14,15^ A compound was classed TdeP positive if it was classified as known (KR) or partial risk (PR) on CredibleMeds which refers to whether there is substantial evidence the drug causes QT prolongation and/or TdeP. A compound is classed as TdeP negative if it was classified as conditional risk (CR), the risk of TdeP is conditional on other factors e.g. drug-drug interaction, or no risk if it wasn’t listed (NR) as was done by Kramer *et al.*^16^ All data is provided in supplemental material.

### Model input data

The percentage block against a given ion-channel inputted into all models was calculated based on the effective therapeutic concentration (EFTPC), which was provided in the original articles, using a pore block model,

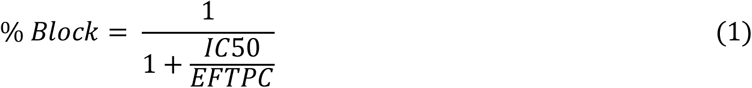

## Models

### Single cell cardiac model simulations

The AP predict platform^25^ was used to simulate the gold-standard and cardiac safety simulator models in all cases except for one simulation study. A MATLAB version of the *gold-standard* model available on the Rudylab website (http://rudylab.wustl.edu) was used when simulating the block of 7 ion-channels since that model on AP predict does not allow blocking of INaL – a current measured in the Crumb *et al.* data-set. The default settings within the AP predict platform were used i.e. 1Hz pacing for 5 minutes with the APD90, time taken for the action potential to repolarise by 90%, recorded using the last cycle. The same protocol was applied in MATLAB when exploring the 7 ion-channels within the O’Hara model i.e. 1Hz pacing for 5 minutes with APD90 recorded using the last cycle. In all simulations drug block was initiated at the beginning of simulations.

### B_net_

We define the difference in block between repolarisation and depolarisation ion-channels as *B_net_*,

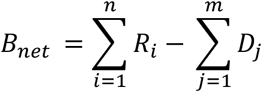

where *R_i_* and *D_j_* represent the percentage block against repolarisation and depolarisation ion-channels respectively for a specific drug.

### Classification evaluation

For each compound the percentage change in APD90 compared to control (no block) from the biophysical model simulations was recorded as was the *B_net_* value. These values were then placed within a logistic regression analysis to assess their correlative value to TdeP risk. This was done via a leave one out cross validation (LOOCV). This involves training a classifier to *n-1* compounds and testing on the *n^th^*. Thus all compounds perform part of the test-set. The predicted probability of risk for each test compound is then used to generate a ROC AUC (area under the receiver operating characteristic curve) and is reported as was done previously.^26^

## Results

### Data

The total number of compounds that are TdeP positive (CredibleMeds known (KR) or partial (PR) risk) versus TdeP negative (CredibleMeds conditional risk (CR) or not listed (NR)) across the 3 datasets of interest can be seen in Figure 1. Although the total number of compounds differs from one data-set to another the proportions that are KR or PR does not.

**Figure 1.**
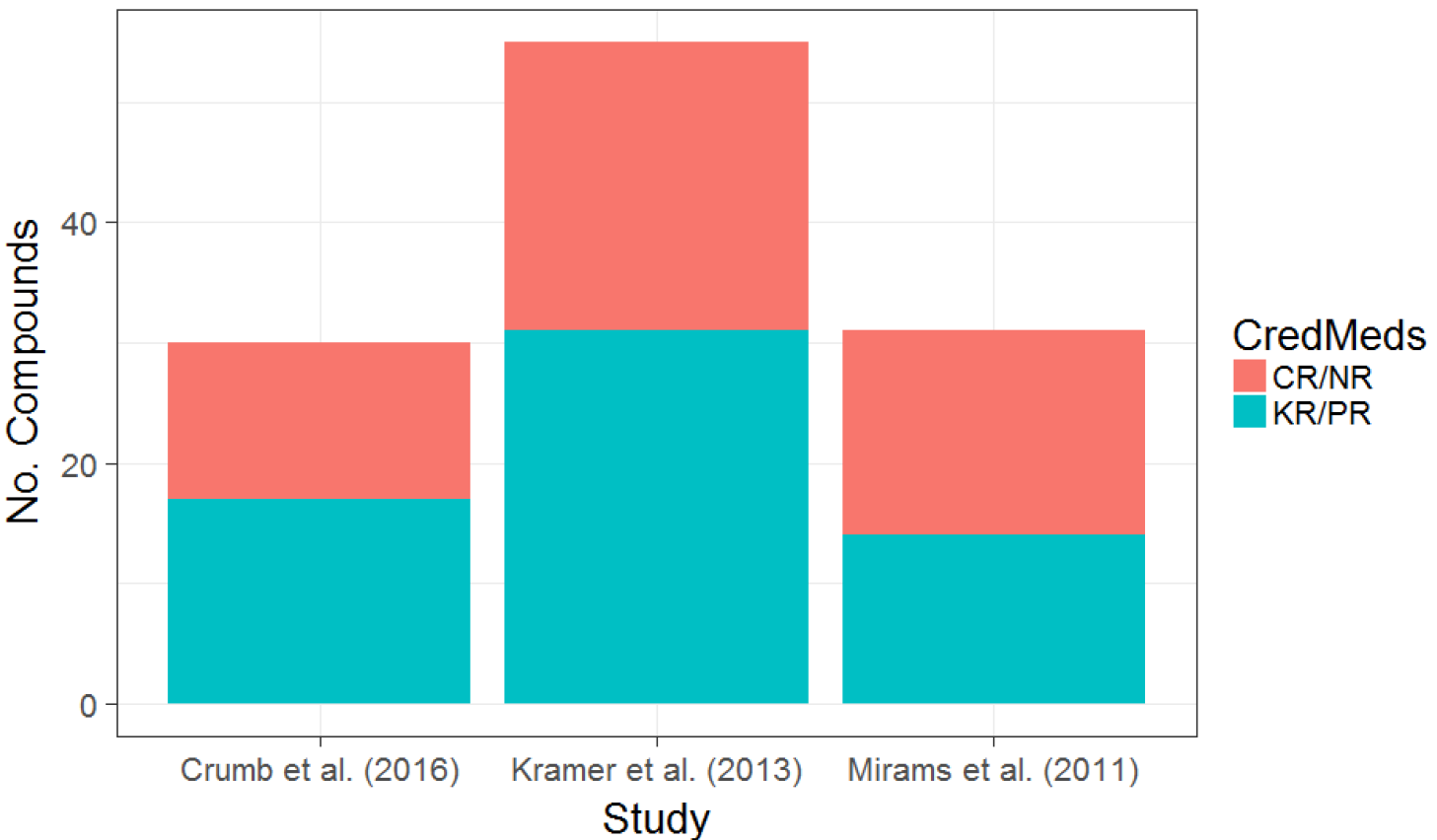
Stacked bar-chart shows the proportion of compounds in each data-set that are TdeP positive (KR/PR on CredMe s database) or TdeP negative (CR on CredMeds database or not listed, NR).

The distribution of block against each ionic current, at the effective therapeutic concentration (EFTPC), across all data-sets can be seen in Figure 2. The plots show that the activity of the compounds is greatest against IKr across all data-sets. After IKr, ICaL appears to be the next channel for which a substantial amount of activity is seen. A somewhat surprising result is the degree of activity against INaL but not INa in the Crumb *et al.* data-set. The amount of activity against INaL in that data-set mirrors that of ICaL activity.

**Figure 2.**
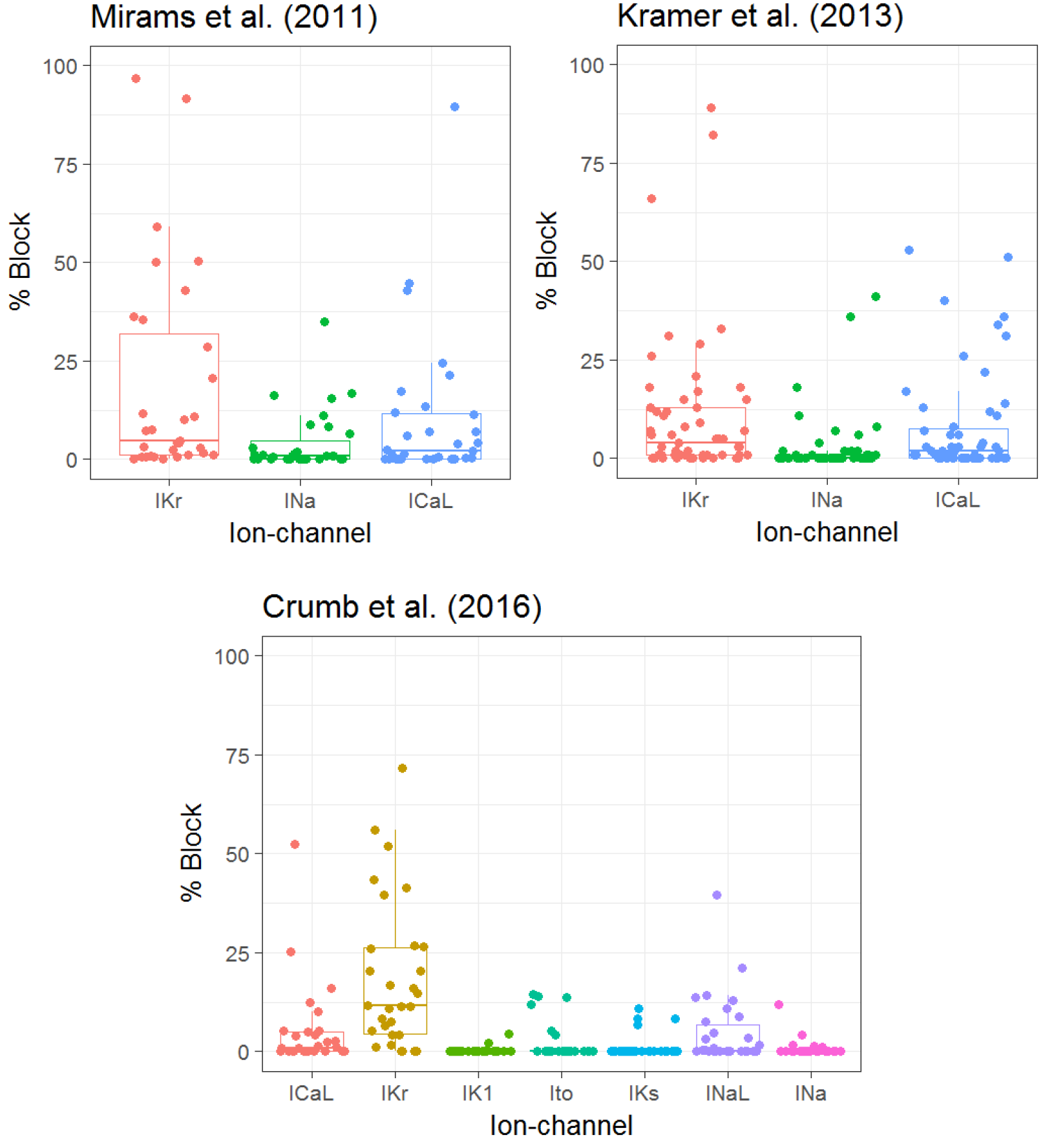
Boxplots show the distribution of block for each ionic current across all 3 data-sets.

### Classification Evaluation

The results of the leave-one-out cross validation for each data-set using various models can be seen graphically in Figure 3 and also in Table 1. For the Mirams *et al.* data-set it’s noticeable that the *gold-standard* model performs no better than using just block against hERG alone neither of which are better than random chance. Both the *cardiac safety simulator* and *B_net_* show a similar improvement over using just hERG block.

**Figure 3.**
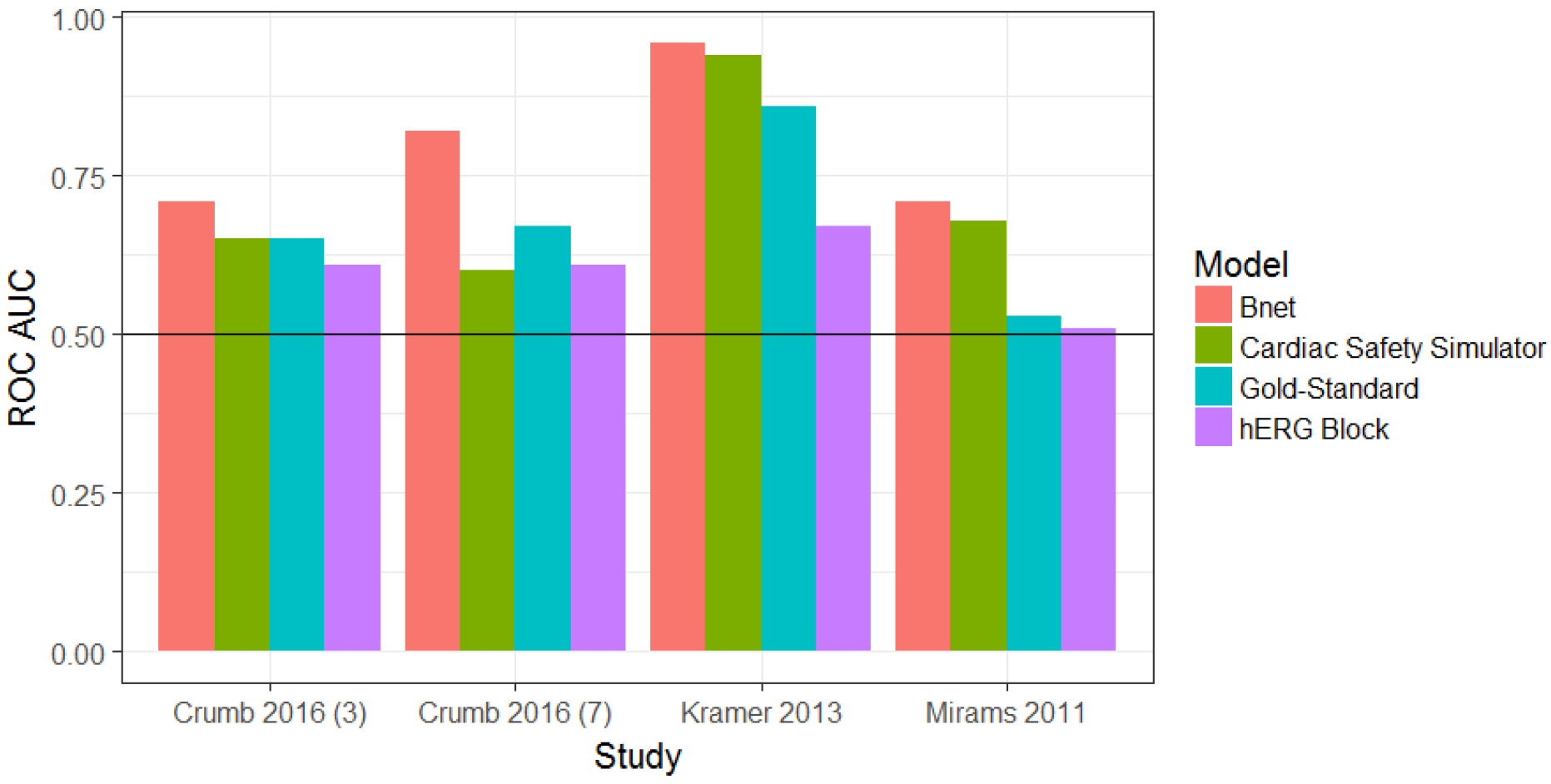
Bar-chart showing the performance of each model across all data-sets studied. The number in parentheses for the Crumb data-set refers to the number of ion-channels used 3 v 7.

**Table 1.**
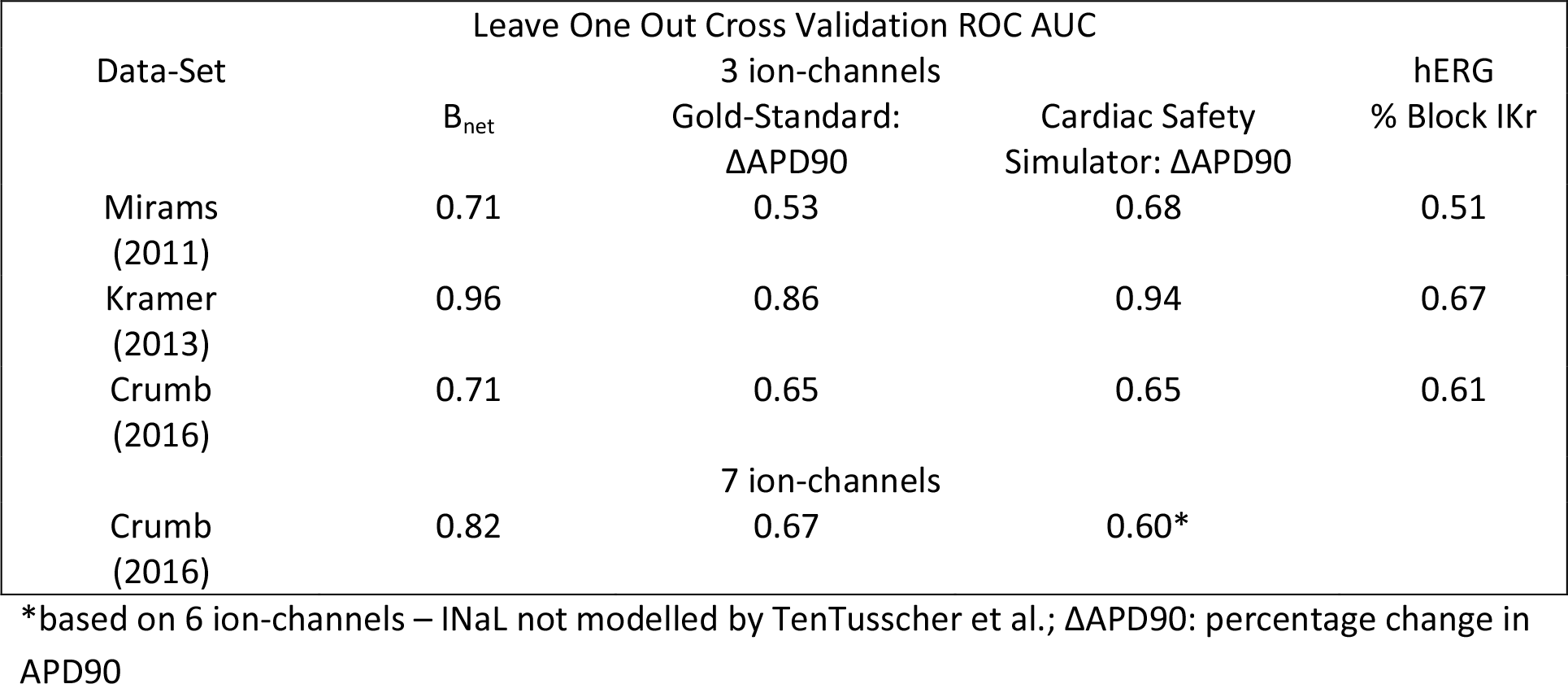
Results of the leave one out cross validation results across all data-sets for all models considered.

Moving onto the Kramer *et al.* data-set the performance of all models improves dramatically over the Mirams *et al.* data-set. Here all 3 models show superior performance over just hERG block. Note that again the *gold-standard* model performance is not as high as *B_net_* or the *cardiac safety simulator*. In addition the difference between *B_net_* and the *cardiac safety simulator* is negligible.

Within the latest data-set by Crumb *et al.* the performance of all models, when using only 3 ion-channels, drops to a level similar to that seen within the Mirams *et al.* data-set. The key difference between the results between those two data-sets is that the *gold-standard* model now shows similar performance to the *cardiac safety simulator*. Furthermore neither biophysical model performs overly better than using hERG block. *B_net_* however appears to give reasonable performance again and appears to show an improvement over using hERG block. Finally if we move onto using all the ion-channel data from the Crumb *et al.* data-set the difference in performance between the models is quite striking. *B_net_*’s performance improves with the addition of more information whereas there is little improvement in either biophysical model. The cardiac safety simulator may even have regressed slightly.

In summary the results show that the performance of the models is data-set dependent. However, within each data-set the *B_net_* model performs just as well if not better than leading biophysical models.

## Discussion

There appears to be a strong belief within the field of ion-channel cardiac drug toxicity that large scale cardiac models are required to answer a well-defined question^10^: can high-throughput ion-channels screening data predict the torsaedgenic risk of a drug in man? The evidence base, that suggests that large-scale models perform better than simpler models for this question, simply does not exist. As previous studies have shown that the performance of the large-scale cardiac models can be mirrored by simpler models.^5^ Furthermore, the simpler model may have potential to outperform large-scale cardiac models.

There were two major caveats in those previous studies. The first relates to the definition of torsadegenic risk, different databases were used, which has been debated within the literature.^19^ Within this study the classification was based on information from CredibleMeds.^14^ Their classification is based on an extensive search of both the literature and public databases and are well known to the clinical community. Another advantage of the CredibleMeds classification is that they do not have a vested interest in the application of mathematical models within drug development.

The second caveat relates to the dimensionality of the ion-channel space, only 3 ion-channels were considered in previous studies.^16,17^ Therefore an understanding as to how generalizable the inferences were on those data-sets to larger dimensions was unknown. This caveat was addressed here by considering a data set by Crumb *et al.* which measured the drug affinity for 7 ion-channels^18^ in addition to the previous data-sets using only 3 ion-channels.^16,17^

Both of these caveats were addressed within this study. Three models were evaluated against the data-sets: 1) the *gold-standard*^8^ single cell model by O’Hara and Rudy^9^; 2) the single cell model by TenTusscher *et al.*^11^ which is used within the *cardiac safety simulator*^12^; 3) a linear model evaluating the net difference in block between ion-channels involved in repolarising and depolarising the action potential, B_net_.^5^ Each model was assessed via a leave-one-out cross validation. (Note that prospective assessment of models is not possible within this field since this would involve developing compounds with a TdeP risk which can be considered unethical.) In addition to using outputs from the aforementioned models within the classification exercise the amount of block against hERG channel was used as a naïve benchmark.

Overall the results showed that *B_net_* was equal if not superior to the biophysical models. The key findings were as follows. Within the Mirams *et al.* data-set the *gold-standard* model was no better than hERG block neither of which was better than random chance. In the largest data-set, by Kramer *et al.*, all models show good performance and highlighted the benefit of measuring more than hERG. When using information on 7 ion-channels within the Crumb *et al.* data-set the performance of *B_net_* was greater than that of the biophysical models. Both of which showed no improvement in performance when moving from 3 to 7 ion-channels unlike *B_net_*. Furthermore the performance of the biophysical models was not all that superior to using only hERG block. In summary the only model which consistently showed the benefit of measuring more than hERG was *B_net_*.

These results may appear surprising but are not uncommon in prediction problems in other fields.^20,21^ The key reason why complex models are not necessarily more predictive than simpler models is due to model error i.e. error in the structure of the model itself.^3^ The concept of model error has not been discussed at all within the cardiac modelling field. Thus the effect of model error on predictivity is largely unknown, although in other fields it tends to dominate prediction uncertainty.^22,23^

A key caveat of the analysis conducted is that the data-sets used may be too small to understand how large a discrepancy there truly is between the different models. However it is hoped that by continuing to assess new data-sets as they become available that the community will eventually have a comprehensive compound list. Other caveats that relate to the *B_net_* model itself are that it doesn’t consider the kinetics of blocking which has been highlighted as an important factor.^24^ However these studies have been on a small numbers of compounds and so a true assessment of the importance of kinetics cannot be determined from those studies alone. If sufficient evidence regarding the importance of drug kinetics does eventually become available the *B_net_* model can first be adapted in one of two possible ways: i) make its variables time-dependent or ii) introduce a scaling factor which accounts for the type of modulation (e.g. slow versus fast etc.). Thus there is a way to improve the model by considering kinetics of drug block if sufficient evidence suggests this will improve predictive/explanatory power․.

In summary the study conducted here highlights the importance of benchmarking. Furthermore it highlights that simple mechanistic models can not only give similar performance to large-scale mechanistic models but can out-perform them. Finally it is hoped this study highlights that there is more than one solution to a problem and that although the question and quality of data dictates the modelling approach it should not dictate the size of the model.

## Supplementary Data

The data-sets used in the analysis are given below. For each compound in each data-set the % block against an ionic current is given together with the % change in APD90 (time taken for the action potential to repolarise by 90%) for the 2 biophysical models under consideration and the compounds classification within the Credible Meds database.

Note that for the Crumb et al. data-set 4 simulation results are provided for each compound. Ohara3 and TenTusscher3 refer to simulations using information on IKr, INa and ICaL only, whereas OharaAll and TenTusscherAll refer to simulations using information on all ionic currents.

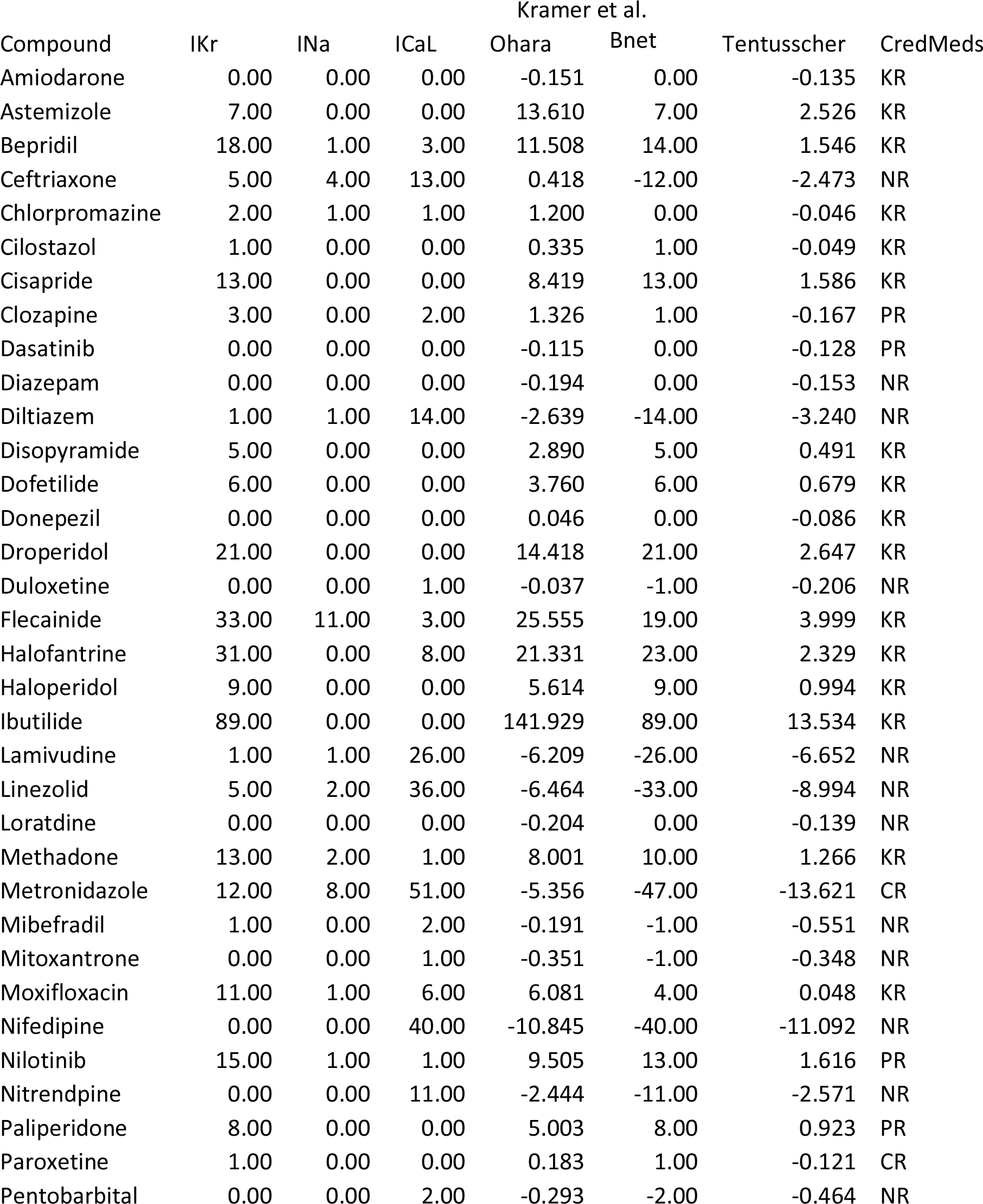

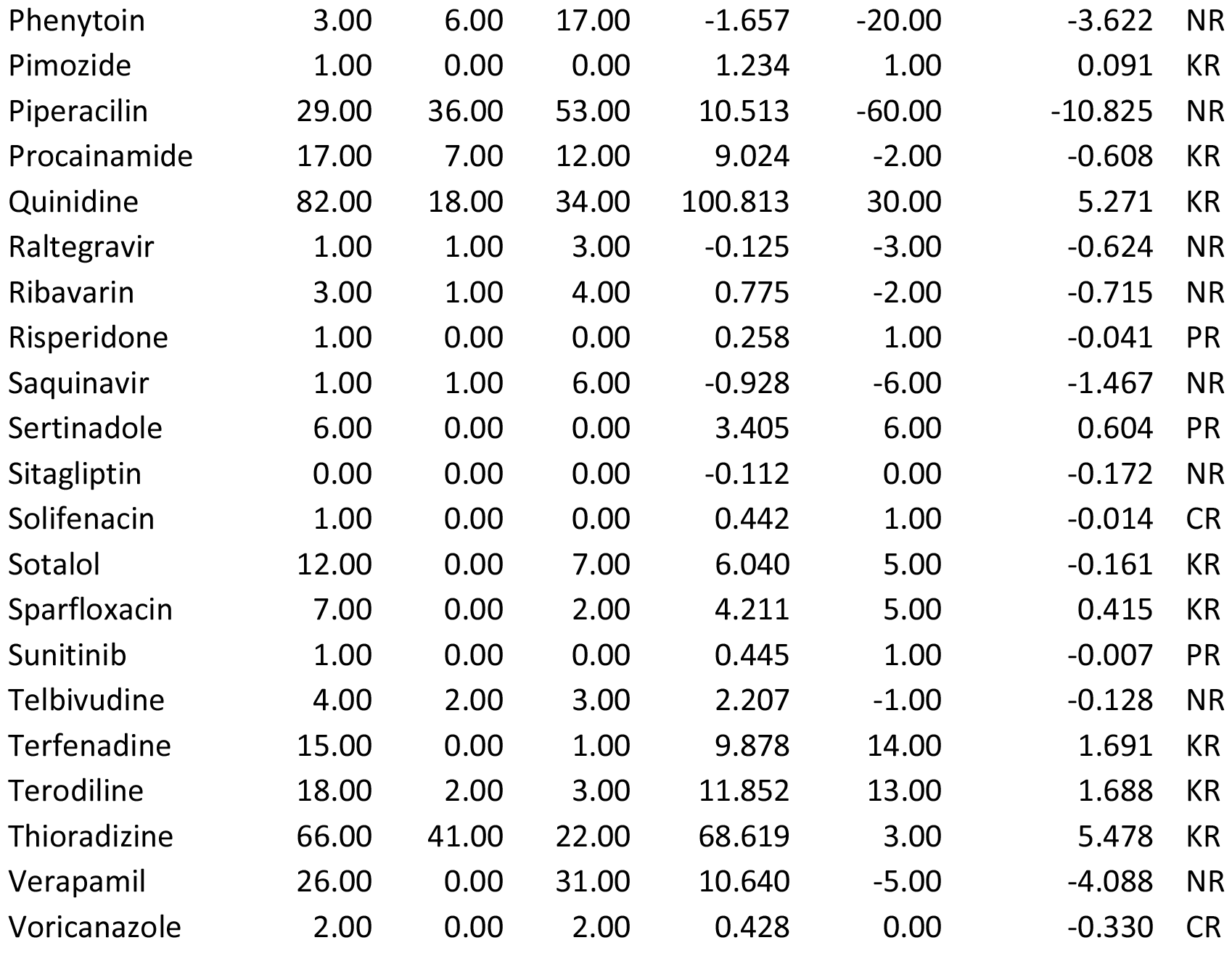

| Compound | IKr | INa | ICaL | Kramer et al. |  |  | CredMeds |
| --- | --- | --- | --- | --- | --- | --- | --- |
|  |  |  |  | Ohara | Bnet | Tentusscher |  |
| Amiodarone | 0.00 | 0.00 | 0.00 | -0.151 | 0.00 | -0.135 | KR |
| Astemizole | 7.00 | 0.00 | 0.00 | 13.610 | 7.00 | 2.526 | KR |
| Bepridil | 18.00 | 1.00 | 3.00 | 11.508 | 14.00 | 1.546 | KR |
| Ceftriaxone | 5.00 | 4.00 | 13.00 | 0.418 | -12.00 | -2.473 | NR |
| Chlorpromazine | 2.00 | 1.00 | 1.00 | 1.200 | 0.00 | -0.046 | KR |
| Cilostazol | 1.00 | 0.00 | 0.00 | 0.335 | 1.00 | -0.049 | KR |
| Cisapride | 13.00 | 0.00 | 0.00 | 8.419 | 13.00 | 1.586 | KR |
| Clozapine | 3.00 | 0.00 | 2.00 | 1.326 | 1.00 | -0.167 | PR |
| Dasatinib | 0.00 | 0.00 | 0.00 | -0.115 | 0.00 | -0.128 | PR |
| Diazepam | 0.00 | 0.00 | 0.00 | -0.194 | 0.00 | -0.153 | NR |
| Diltiazem | 1.00 | 1.00 | 14.00 | -2.639 | -14.00 | -3.240 | NR |
| Disopyramide | 5.00 | 0.00 | 0.00 | 2.890 | 5.00 | 0.491 | KR |
| Dofetilide | 6.00 | 0.00 | 0.00 | 3.760 | 6.00 | 0.679 | KR |
| Donepezil | 0.00 | 0.00 | 0.00 | 0.046 | 0.00 | -0.086 | KR |
| Droperidol | 21.00 | 0.00 | 0.00 | 14.418 | 21.00 | 2.647 | KR |
| Duloxetine | 0.00 | 0.00 | 1.00 | -0.037 | -1.00 | -0.206 | NR |
| Flecainide | 33.00 | 11.00 | 3.00 | 25.555 | 19.00 | 3.999 | KR |
| Halofantrine | 31.00 | 0.00 | 8.00 | 21.331 | 23.00 | 2.329 | KR |
| Haloperidol | 9.00 | 0.00 | 0.00 | 5.614 | 9.00 | 0.994 | KR |
| Ibutilide | 89.00 | 0.00 | 0.00 | 141.929 | 89.00 | 13.534 | KR |
| Lamivudine | 1.00 | 1.00 | 26.00 | -6.209 | -26.00 | -6.652 | NR |
| Linezolid | 5.00 | 2.00 | 36.00 | -6.464 | -33.00 | -8.994 | NR |
| Loratadine | 0.00 | 0.00 | 0.00 | -0.204 | 0.00 | -0.139 | NR |
| Methadone | 13.00 | 2.00 | 1.00 | 8.001 | 10.00 | 1.266 | KR |
| Metronidazole | 12.00 | 8.00 | 51.00 | -5.356 | -47.00 | -13.621 | CR |
| Mibefradil | 1.00 | 0.00 | 2.00 | -0.191 | -1.00 | -0.551 | NR |
| Mitoxantrone | 0.00 | 0.00 | 1.00 | -0.351 | -1.00 | -0.348 | NR |
| Moxifloxacin | 11.00 | 1.00 | 6.00 | 6.081 | 4.00 | 0.048 | KR |
| Nifedipine | 0.00 | 0.00 | 40.00 | -10.845 | -40.00 | -11.092 | NR |
| Nilotinib | 15.00 | 1.00 | 1.00 | 9.505 | 13.00 | 1.616 | PR |
| Nitrendipine | 0.00 | 0.00 | 11.00 | -2.444 | -11.00 | -2.571 | NR |
| Paliperidone | 8.00 | 0.00 | 0.00 | 5.003 | 8.00 | 0.923 | PR |
| Paroxetine | 1.00 | 0.00 | 0.00 | 0.183 | 1.00 | -0.121 | CR |
| Pentobarbital | 0.00 | 0.00 | 2.00 | -0.293 | -2.00 | -0.464 | NR |
| Phenytoin | 3.00 | 6.00 | 17.00 | -1.657 | -20.00 | -3.622 | NR |
| Pimozide | 1.00 | 0.00 | 0.00 | 1.234 | 1.00 | 0.091 | KR |
| Piperacilin | 29.00 | 36.00 | 53.00 | 10.513 | -60.00 | -10.825 | NR |
| Procainamide | 17.00 | 7.00 | 12.00 | 9.024 | -2.00 | -0.608 | KR |
| Quinidine | 82.00 | 18.00 | 34.00 | 100.813 | 30.00 | 5.271 | KR |
| Raltegravir | 1.00 | 1.00 | 3.00 | -0.125 | -3.00 | -0.624 | NR |
| Ribavarin | 3.00 | 1.00 | 4.00 | 0.775 | -2.00 | -0.715 | NR |
| Risperidone | 1.00 | 0.00 | 0.00 | 0.258 | 1.00 | -0.041 | PR |
| Saquinavir | 1.00 | 1.00 | 6.00 | -0.928 | -6.00 | -1.467 | NR |
| Sertinadole | 6.00 | 0.00 | 0.00 | 3.405 | 6.00 | 0.604 | PR |
| Sitagliptin | 0.00 | 0.00 | 0.00 | -0.112 | 0.00 | -0.172 | NR |
| Solifenacin | 1.00 | 0.00 | 0.00 | 0.442 | 1.00 | -0.014 | CR |
| Sotalol | 12.00 | 0.00 | 7.00 | 6.040 | 5.00 | -0.161 | KR |
| Sparfloxacin | 7.00 | 0.00 | 2.00 | 4.211 | 5.00 | 0.415 | KR |
| Sunitinib | 1.00 | 0.00 | 0.00 | 0.445 | 1.00 | -0.007 | PR |
| Telbivudine | 4.00 | 2.00 | 3.00 | 2.207 | -1.00 | -0.128 | NR |
| Terfenadine | 15.00 | 0.00 | 1.00 | 9.878 | 14.00 | 1.691 | KR |
| Terodiline | 18.00 | 2.00 | 3.00 | 11.852 | 13.00 | 1.688 | KR |
| Thioradizine | 66.00 | 41.00 | 22.00 | 68.619 | 3.00 | 5.478 | KR |
| Verapamil | 26.00 | 0.00 | 31.00 | 10.640 | -5.00 | -4.088 | NR |
| Voriconazole | 2.00 | 0.00 | 2.00 | 0.428 | 0.00 | -0.330 | CR |

| Compound | Mirams et al. |  |  |  |  |  |  |
| --- | --- | --- | --- | --- | --- | --- | --- |
|  | INa | ICaL | IKr | Bnet | Ohara | Tentusscher | CredMeds |
| Ajmaline | 15.46 | 2.07 | 59.06 | 41.52 | 57.285 | 8.149 | NR |
| Amiodarone | 0.01 | 0.18 | 1.64 | 1.44 | 1.738 | 0.203 | KR |
| Amitriptyline | 0.20 | 0.35 | 1.23 | 0.68 | 0.495 | -0.054 | CR |
| Bepridil | 0.88 | 13.52 | 50.00 | 35.59 | 40.320 | 4.083 | KR |
| Chlorpromazine | 0.88 | 0.00 | 2.52 | 1.64 | 1.405 | 0.197 | KR |
| Cibenzoline | 11.12 | 24.55 | 4.14 | -31.53 | 2.484 | -0.177 | NR |
| Cisapride | 0.03 | 0.00 | 42.98 | 42.95 | 35.008 | 5.872 | KR |
| Desipramine | 6.63 | 5.94 | 7.21 | -5.37 | 3.720 | -0.439 | PR |
| Diltiazem | 1.34 | 21.33 | 0.70 | -21.97 | -4.787 | -5.206 | NR |
| Diphenhydramine | 0.08 | 0.01 | 0.65 | 0.55 | 0.189 | -0.058 | CR |
| Dofetilide | 0.00 | 0.00 | 28.57 | 28.57 | 20.757 | 3.752 | KR |
| Fluvoxamine | 0.95 | 7.14 | 10.84 | 2.75 | -0.823 | -1.581 | CR |
| Haloperidol | 0.05 | 0.21 | 11.76 | 11.50 | 7.482 | 1.373 | KR |
| Imipramine | 2.86 | 1.26 | 3.02 | -1.10 | 1.618 | 0.008 | PR |
| Mexiletine | 8.76 | 3.97 | 7.63 | -5.10 | 4.510 | 0.082 | NR |
| Mibefradil | 1.21 | 7.14 | 0.66 | -7.69 | -1.133 | -1.636 | NR |
| Nifedipine | 0.02 | 11.37 | 0.00 | -11.39 | -2.609 | -2.728 | NR |
| Nitrendipine | 0.01 | 89.61 | 0.03 | -89.59 | -24.736 | -37.183 | NR |
| Phenytoin | 8.41 | 4.19 | 4.31 | -8.29 | 2.252 | -0.415 | NR |
| Pimozide | 1.82 | 0.61 | 4.76 | 2.33 | 2.792 | 0.373 | KR |
| Prenylamine | 0.67 | 1.35 | 20.73 | 18.71 | 14.000 | 2.366 | NR |
| Propafenone | 16.84 | 11.81 | 35.39 | 6.74 | 26.263 | 2.362 | NR |
| Propranolol | 1.22 | 0.14 | 0.91 | -0.46 | 0.391 | -0.041 | NR |
| Quetiapine | 0.19 | 0.32 | 0.57 | 0.05 | 0.090 | -0.133 | CR |
| Quinidine | 16.32 | 17.18 | 91.52 | 58.02 | 154.967 | 10.943 | KR |
| Risperidone | 0.00 | 0.00 | 1.19 | 1.19 | 0.520 | 0.015 | PR |
| Sertindole | 0.07 | 0.02 | 10.20 | 10.11 | 6.428 | 1.204 | PR |
| Tedisamil | 0.42 | 0.00 | 3.29 | 2.86 | 1.865 | 0.293 | NR |
| Terfenadine | 0.92 | 2.34 | 50.28 | 47.02 | 43.313 | 6.529 | KR |
| Thioridazine | 34.85 | 42.96 | 96.74 | 18.93 | 185.571 | 6.492 | KR |
| Verapamil | 0.19 | 44.75 | 36.16 | -8.79 | 16.067 | -6.865 | NR |

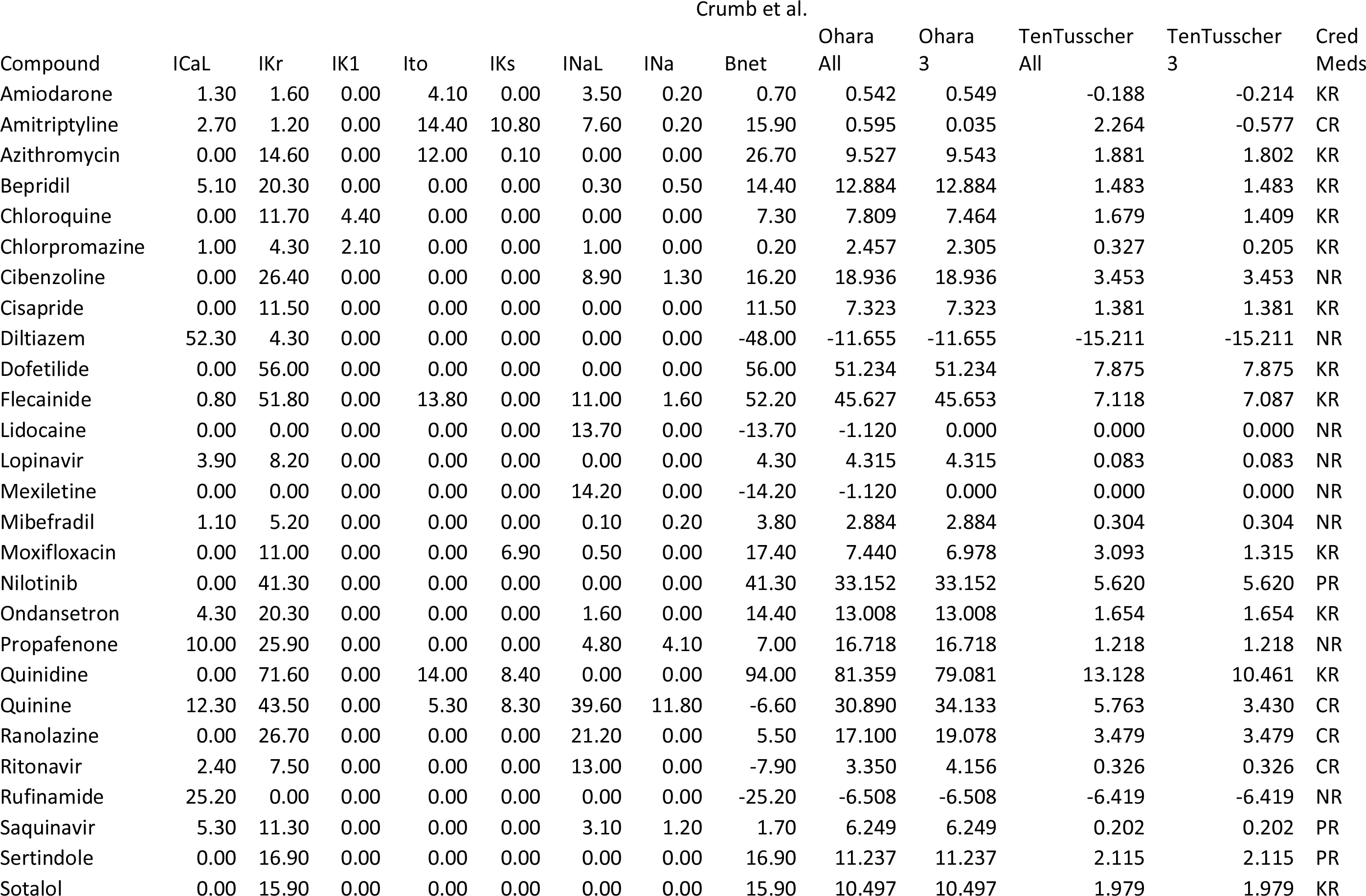

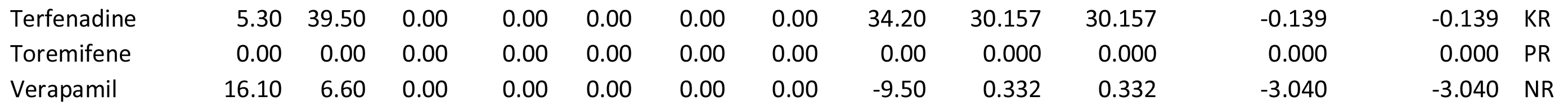

